# Magnetic Tweezers Experiments Reveal Increased Mechanical Sensitivity of the F2561Y Mutant in Von Willebrand Factor

**DOI:** 10.64898/2026.06.05.730321

**Authors:** Caroline Körösy, Stefanie D. Pritzl, Tobias Obser, Maria Brehm, Jan Lipfert

## Abstract

The large glycoprotein von Willebrand factor (VWF) is essential for primary hemostasis. By sensing hydrodynamic forces in the bloodstream, VWF elongates under elevated shear, increasing its adhesiveness to platelets. This mechanosensitive behavior arises from its multidomain architecture, where distinct structural elements respond at different force scales. The stem, formed by interactions between C-domains of adjacent monomers in VWF dimers, opens at relatively low forces (~1 pN), representing an early step in force activation. The F2561Y mutation in the C4 domain is a gain-of-function mutation associated with increased shear sensitivity and myocardial infarction, yet its effect on VWF mechanics in the physiologically relevant low-force regime remains unresolved. Here, we combine single-molecule magnetic tweezers with AFM imaging to investigate how F2561Y alters VWF stem dynamics. Magnetic tweezers measurements reveal that A2 domain unfolding remains unchanged in the mutant, whereas stem opening transitions occur at significantly lower forces compared to wildtype. The midpoint force for stem opening is reduced by approximately one order of magnitude, indicating a pronounced destabilization of the compact stem conformation. AFM imaging independently confirms a higher population of open stem states in the mutant, even without applied force. Our findings demonstrate that F2561Y selectively destabilizes the VWF stem and increases its force sensitivity without perturbing global domain stability, providing a direct molecular explanation for its enhanced shear responsiveness and prothrombotic phenotype. More broadly, our work establishes magnetic tweezers as a powerful approach to resolve low-force conformational equilibria in mechanosensitive proteins and to dissect the mechanical impact of disease-associated mutations.

**Highlights:**

- The F2561Y mutation lowers the force required to open the VWF stem.
- F2561Y mutant VWF dimers exhibit more open conformations compared to wildtype.
- Force sensitivity provides a mechanism for the prothrombic phenotype of F2561Y.

## Introduction

Von Willebrand factor (VWF) is a large plasma glycoprotein, critically involved in primary hemostasis. VWF circulates in the blood stream in the form of long, multimeric chains, which under normal conditions adopt compact, globular conformations. Mature VWF monomers consist of the D′-D3-A1-A2-A3-D4-C1-C2-C3-C4-C5-C6-CK domains [1, 2] (Figure 1A). In the circulating protein, pairs of monomers are covalently linked *via* C-terminal disulfide bonds within their cysteine-knot (CK) domains to form VWF dimers, which constitute the smallest repeating unit in VWF multimers, and are further connected by N-terminal disulfide bonds to generate long VWF concatamers.

**Figure 1:**
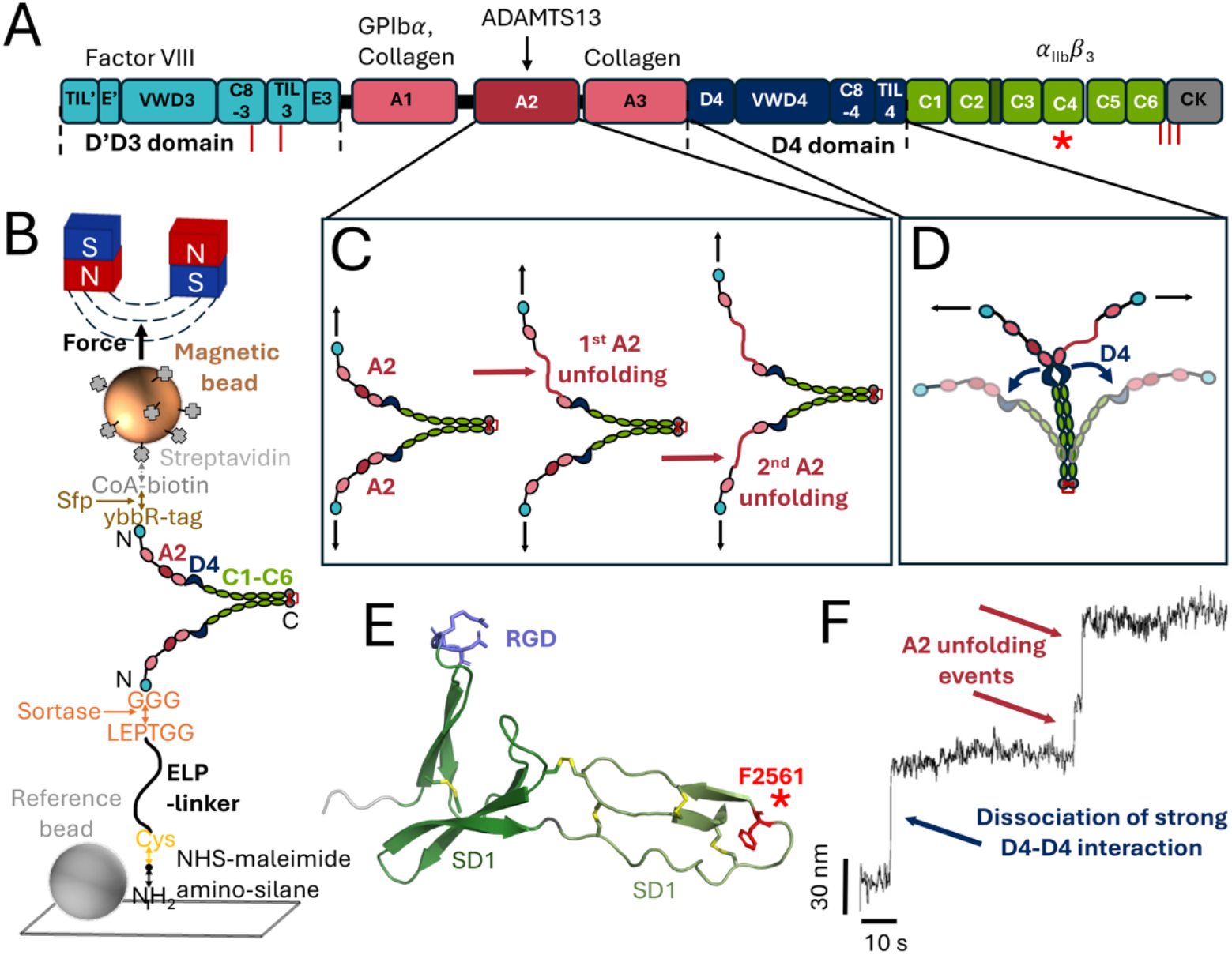
VWF domain architecture and magnetic tweezers assay. A) Domain organization of a mature VWF monomer. Binding sites of key interaction partners of VWF are indicated on top, cysteine residues involved in C- and N-terminal disulfide bond formation for dimerization and multimerization, respectively, are highlighted in dark red below, and the mutation site in of F2561Y in the C4 domain is indicated with a red asterisk. B) Attachment strategy for VWF dimers in the magnetic tweezers setup. The construct is N-terminally tethered to an amino-functionalized glass surface *via* an elastin-like polypeptide spacer using sortase-mediated coupling and to a streptavidin-coated superparamagnetic bead *via* biotin-streptavidin interaction. Surface-immobilized polystyrene beads are used as a reference for drift correction. External magnets exert force on the bead and the attached protein dimer. C) Schematic representation of independent unfolding of the two A2 domains under force, producing a characteristic two-step signature that serves as a molecular fingerprint. D) Schematic illustration of the dissociation of the D4-D4 interaction that occurs in some, but not all traces and gives rise to an additional stepwise unfolding pattern. E) Cartoon representation of the VWF C4 domain based on its NMR structure [31] (PDB 6FWV). Subdomains are shown in dark green (SD1) and light green (SD2), with disulfide bonds highlighted in yellow. The integrin-binding motif (RGD) is shown as blue sticks, and mutation site (residue 2561) is indicated in red, showing the wildtype. F) Representative extension-time trace recorded at 12.5 pN showing sequential unfolding of individual A2 domains and dissociation of the D4-D4 interaction in a single VWF dimer.

Force-induced structural changes are central to VWF regulation [3-6] and the tightly regulated mechanosensitive behavior of VWF arises from the interplay between VWF’s molecular architecture and the forces encountered in the vasculature. Due to their concatameric, multidomain organization VWF multimers are activated and undergo pronounced conformational changes upon sensing elevated hydrodynamic forces, as they occur at sites of vascular injury [3, 4]. Importantly, the peak shear forces acting on the multimer increase with the square of its effective length [4, 7, 8]. Therefore, conformational changes that result in elongation increase the acting forces, which set off additional transitions, giving rise to a cascade of force-induced molecular events. Within this hierarchical mechanical response, dissociation of interdomain interactions can expose structural elements that respond at different force scales. Pairwise interactions among the C-domains form the VWF stem, which can reversibly open and close in a zipper-like manner and has been shown to respond to low forces in the range of ~1 pN [9]. However, the VWF stem can be stabilized in the compact conformation by D4-D4 interdomain interaction, which shields the stem from opening in approximately half of the VWF dimers [10]. Once the stabilizing D4–D4 interaction is released, the dimeric stem can undergo reversible opening and closing, thereby contributing to the early conformational response of VWF. The A2 domain serves as a mechanical sensor at higher forces: lacking long-range disulfide bonds, it unfolds under tensile force in the range of 10-20 pN [8, 9, 11], exposing a cryptic cleavage site for the metalloprotease ADAMTS13, thereby controlling multimer size and acivity. This intrinsic length-dependent foce amplification enables VWF to function as a flow sensor, selectively acting under elevated shear conditions associated with vascular injury or disturbed flow. Through force-induced structural changes, VWF exposes binding sites for platelet receptors. Under physiological condition, this enables platelet recruitment at sites of vascular injury and formation of a hemostatic plug [3]. However, dysregulated force responsiveness of VWF can result in undesirable blood clotting and increase the risk of arterial thrombosis [12, 13].

Mutations in the *VWF* gene cause von Willebrand Disease (VWD) [14, 15] by affecting the quantity (Types 1,3) or quality (Type 2: 2A, 2B, 2M, 2N) of VWF, leading to prolonged bleeding [16, 17]. VWD is the most common inherited bleeding disorder, affecting roughly 1% of the population and hundreds of mutations exist, causing varied defects ranging from reduced VWF levels (Type 1) to complete absence (Type 3) or specific functional impairments (Type 2) [16, 18-21].

Gain-of-function mutations in VWF increase its activity and can lead to divergent clinical phenotypes. In VWD Type 2B, such mutations, most commonly affecting the A1 domain, enhance VWF binding affinity for platelet glycoprotein Ibα, causing spontaneous platelet-VWF interactions, accelerated platelet clearance and thrombocytopenia, ultimately impairing effective hemostasis and causing bleeding symptoms [22-25]. In contrast, gain-of-function mutations affecting domains involved in VWF conformational regulation, including the C4 domain, can increase structural flexibility and sensitivity to shear stress, promoting excessive activation under flow [26, 27]. This hyperadhesive VWF promotes platelet aggregation and increases the risk of pathological arterial thrombosis. Thus, depending on the affected domain and the specific biophysical consequences, VWF gain-of-function mutations can lead to either bleeding due to platelet depletion or thrombotic complications due to excessive platelet adhesion.

The F2561Y variant of VWF is a gain-of-function mutation with prothrombotic properties [26] located within the C4 domain, which harbors the RGD motif for platelet integrin GPIIb/IIIa [28-30]. Carriers of the Y2561 allele exhibit an increased risk of recurrent myocardial infarction before the age of 55 years, particularly among women, suggesting an elevated prothrombotic risk [26]. Functional studies demonstrate increased shear sensitivity, reduced critical shear rates for platelet aggregate formation, and enhanced aggregate size compared to wildtype VWF [26, 27]. Circular dichroism spectroscopy indicates preserved folding of isolated C domains, but altered spectra in dimeric F2561Y constructs [26] suggest perturbations of interdomain organization rather than intrinsic instability. NMR spectroscopy reveals no major structural changes within the VWF F2561Y C4 domain or its integrin-binding capability [31], which supports the hypothesis that the gain-of-function phenotype arises from altered spatial organization of the C4 domain within the neighboring stem domains. AlphaFold structural models predict weakened dimeric interface interactions in the F2561Y variant, favoring more open stem conformations [27]. Optical tweezers experiments further demonstrate altered stem dynamics and reduced events of dimeric D4 interaction disruption, supported by AFM images, revealing a significant shift toward open stem configurations in the mutant protein [27]. However, the direct impact of F2561Y on stem mechanics under physiologically relevant forces remains unresolved, since the low-force regime (≤ 1 pN), critical for VWF stem opening [9], is technically challenging to access using single-molecule force spectroscopy assays [32, 33].

Here we use single-molecule magnetic tweezers (MT), which enable stable force application with sub-piconewton resolution over extended time scales while simultaneously monitoring multiple molecules in parallel [9, 33, 34]. The MT approach allows us to resolve reversible conformational equilibria governing stem opening in the physiologically relevant regime of low forces. Using MT, we directly compare stem mechanics of wildtype and F2561Y VWF dimers. We show that A2 domain unfolding remains unaffected by the F2561Y mutation, whereas stem opening occurs at significantly lower forces and stem transitions are observed more frequently in the mutant. Our results directly demonstrate that F2561Y lowers the mechanical threshold for stem opening and activation.

To complement the single-molecule force manipulation measurements, we analyzed VWF dimers by AFM imaging in the absence of applied force. Our AFM imaging data directly demonstrate a statistically significant larger fraction of dimers in open stem conformation for F2561Y compared to wildtype, in agreement with previous qualitative results from AFM imaging [27] and fully consistent with our magnetic tweezers results.

Together, our findings demonstrate that the F2561Y mutation enhances stem opening at low forces, providing a direct mechanistic link between altered stem dynamics and the prothrombotic phenotype, and establishing magnetic tweezers as a powerful approach to resolve such conformational transitions in the physiologically relevant low-force regime.

## Materials and Methods

### VWF Constructs

For attachment and measurements in MTs, we use hetero-bifunctional VWF dimers, modified at the two N-termini, at one end with an ybbR-tag (DSLEFIASKLA) [35] for later biotinylation, and at the other terminus with a twin-Strep-tag II (WSHPQFEKGGGSGGGSGGGSWSHPQFEK) for purification, followed by a TEV protease cleavage site to remove the strep-tag after purification, and the sortase recognition motif *GG* for site-specific immobilization. We use mature dimeric constructs, which lack the VWF pro-peptide (domains D1 and D2) in both monomers. For the mutant F2561Y, phenylalanine at position 2561 is exchanged to tyrosine.

The plasmids were constructed and the proteins expressed as previously described [9, 10, 26]. The dimeric constructs were further purified *via* a StrepTrap affinity chromatography column (Citiva) on an ÄKTA Explorer FPLC system (GE Healthcare) in VWF measurement buffer (Supplementary Table S1) and eluted with additional 2.5 mM d-desthiobiotin (Sigma-Aldrich). The eluates were concentrated in VWF measurement buffer using Amicon Ultra MWCO 100 kDa centrifugal filters (Merck). AFM imaging of the purified constructs confirmed structurally intact VWF dimers for further use in MTs.

### Flow Cell Preparation and Protein Attachment

We assemble flow cells from two glass coverslips (#1, 24 mm × 60 mm; Carl Roth, Karlsruhe, Germany) separated by a single layer of parafilm with a cutout to form a flow channel. Bottom coverslips are functionalized with ELP linkers (Supplementary Materials and Methods and Refs. [36-38]) for protein attachment following the protocol by Ott *et al*. [37]. Cleaned slides were silanized with 3(-aminopropyl)triethoxysilane (Thermo Scientific) and coated with a sulfosuccinimidyl 4-(n-maleimidomethyl)cyclohexane-1-carboxylate cross-linker (Thermo Scientific) in 50 mM HEPES, pH 7.5. ELP Linkers [9] (in 50 mM sodium phosphate, 50 mM NaCl, 10 mM EDTA, pH 7.2) were linked to the thiol-reactive maleimide groups *via* an N-terminal cysteine and remaining unreacted maleimide groups were saturated by adding 10 mM L-cysteine. Each functionalization step was followed by extensive rinsing with ultrapure water. Finally, polystyrene beads of 3 µm diameter (Polybead Microspheres) in ethanol were immobilized on the surface to serve as reference beads for drift correction in the later measurement.

The top coverslips were prepared with two holes of ~1 mm diameter for fluid inlet and outlet. Mounted flow cells were incubated with 1% (w/v) caseine (Sigma-Aldrich) for 1 h, followed by rinsing with ~ 1 ml of VWF measurement buffer (Supplementary Table S1) for passivation.

Prior to the measurements, CoA-biotin (NEB) was coupled to the ybbR-tag of the VWF constructs in a bulk reaction of 20 µl in measurement buffer with 5 µM sfp-phosphopantetheinyl transferase (Supplementary Materials and Methods and Refs.[37]) for 1 h at 37 ^°^C. For cleavage of the strep-tag at one VWF-N-terminus at the TEV site, desalted TEV protease (NEB) was added to a final concentration of 25 µM and incubated for 2 h at room temperature. The biotinylated and activated protein was diluted to ~ 20 nM in measurement buffer and incubated in the flow cell with 1 µM eSortaseA (Supplementary Materials and Methods and Refs. [39, 40]) for 30 min for covalent linkage to the immobilized ELP linkers on the glass surface, subsequently flushed with 1 ml VWF measurement buffer (Supplementary Table S1).

Streptavidin-coated superparamagnetic beads of 2.8 µM diameter (M270, Invitrogen) in measurement buffer were flushed into the flow cell for attachment to the proteins *via* non-covalent biotin-streptavidin interactions. Unbound magnetic beads were flushed out with measurement buffer. Measurements were performed at room temperature on a custom-built MT setup (Supplementary Materials and Methods and Refs. [9, 34, 41-47]).

## Data Analysis

Data were analyzed using custom-written Matlab and python scripts. From the measurements we obtain extension *vs*. time traces, from which we subtract the position of a reference bead to correct for drift. Dwell times and extensions increments for A2-domain unfolding events were determined from extension time traces (Supplementary Figure S2).

To analyze transitions in the VWF stem, we apply a locally weighted non-parametric regression (LOESS) filter to smooth the extension-time traces and reduce noise. A midpoint-extension threshold was then set as the midpoint between the maximum and minimum extension of the smoothed signal to distinguish between states. Raw data points were subsequently classified as belonging to closed (below threshold) or open (above threshold) stem conformation. The resulting fractions were fit with a two-state model assuming an Arrhenius-like force dependence [9, 48] for each analyzed molecule individually:

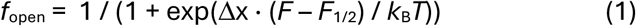

where *k*_B_ is the Boltzmann constant, T the absolute temperature, and *F*_1/2_ and Δx are fitting parameters that represent the midpoint force and a length-scale parameter characterizing the cooperativity of the transition, respectively.

### AFM Imaging

For AFM imaging, we deposited 20 µl of VWF wildtype or F2561Y mutant dimer constructs at nanomolar concentration in VWF measurement buffer (Supplementary Table S1) on 0.1% PLL-coated (0.1%) mica and incubated for 2 min. The samples were rinsed with ultrapure water and blowdried in a gentle stream of nitrogen gas [49].

Images were obtained using a Nanowizard Ultraspeed 2 high speed AFM (JPK) with triangular FASTSCAN A probes (Bruker) with a driving frequency of ~1400 kHz and a spring constant of 18 N/m. The samples were scanned in tapping mode in air with a resolution of ~1 nm/pixel and a line scanning speed of 3 Hz. AFM images were processed using the MountainsSPIP 10 analysis software (Digital Surf) applying line-by-line correction and polynomial background subtraction. Only clearly identifiable, isolated single molecules were considered for analysis. Molecules were classified as adopting a closed stem conformation when they exhibited three distinct protrusions: two flanks of similar length and thickness, corresponding to the folded D’D3 domains, and one longer, thicker flank corresponding to the compacted stem region,with the flanks being connected by a higher feature, attributed to the closed D4 domains. Dimers were classified as being in an open conformation when they displayed two extended flanks with a combined length consistent with the D’D3, D4 and C domains. In the open conformations, two distinct higher features were observed, corresponding to the folded D’D3 and D4 domains.

## Results

We use magnetic tweezers to investigate structural transitions under force in von Willebrand factor at the single molecule level. Dimeric VWF constructs are tethered between a glass surface and superparamagnetic beads in a flow cell (Figure 1B, Methods), mimicking how VWF dimer would be exposed to force in VWF multimers under shear flow. Surface attachment is achieved using our elastin-like polypeptide linker approach (Supplementary Materials and Methods) that enables stable attachment while minimizing surface interactions [9, 36, 37]. A pair of external magnets exerts precisely calibrated stretching forces on the beads and thereby to the attached proteins. We compare force-induced conformational transition in wildtype VWF dimers with those of the clinical gain-of-function mutant F2561Y, which carries a point mutation in the C4 domain of the stem. All measurements were performed in buffer at pH 7.4 containing 1 mM Mg^2+^ and Ca^2+^, representative of physiological blood plasma conditions.

Using our magnetic tweezers force spectroscopy assay, we investigate several key transitions in wildtype and F2561F mutant VWF. First, we observe unfolding of the A2 domains, which provides a unique fingerprint to identify single and correctly attached VWF dimers (Figure 1C,F). Second, the assay can induce the dissociation of the strong interaction of D4 domains, which initially shields the VWF stem from external forces in a fraction of VWF dimers (Figure 1D,F). Finally, we focus on the reversible opening and closing transitions of the VWF stem and identify how they are altered by the F2561F mutation.

### A2 Domain Unfolding is not Affected by the F2561Y Mutation

The VWF A2 domain lacks long-range disulfide bonds to protect it from unfolding by tensile force [8, 11]. This structural feature allows the A2 domain to function as a force sensor, enabling it to unfold and expose its buried ADAMTS13 cleavage site in response to shear force [50]. The two A2 domains in the VWF dimer give rise to a characteristic two-step unfolding pattern in our assay that we use as a reliable molecular fingerprint for identifying correctly tethered VWF dimers (Figure 2A, Supplementary Figure S2) [9, 51]. Comparison of wildtype and F2561Y VWF dimers reveals indistinguishable A2 unfolding behavior, with similar unfolding rates: At 12.5 pN, the mean of the dwell time (mean ± SD) is 33.6 ± 26.7 s for wildtype and 36.7 ± 23.9 s for F2561Y, with no significant difference (Welch’s *t*-test: *p* = 0.71). Similarly, the length increases per unfolding event are 32.5 ± 7.7 nm for wildtype and 35.7 ± 8.6 nm for F2561Y, again identical within error (Welch’s *t*-test: *p* = 0.24) (Figure 2B,C). Both the unfolding rate and length increases are consistent with previously reported values [9]. Our results indicate that the F2561Y mutation does not affect the mechanical unfolding properties of the A2 domains. This is consistent with the spatial separation of A2 from the C4 domain within the stem, suggesting that the mutation does not globally destabilize VWF but instead exerts localized effects.

**Figure 2:**
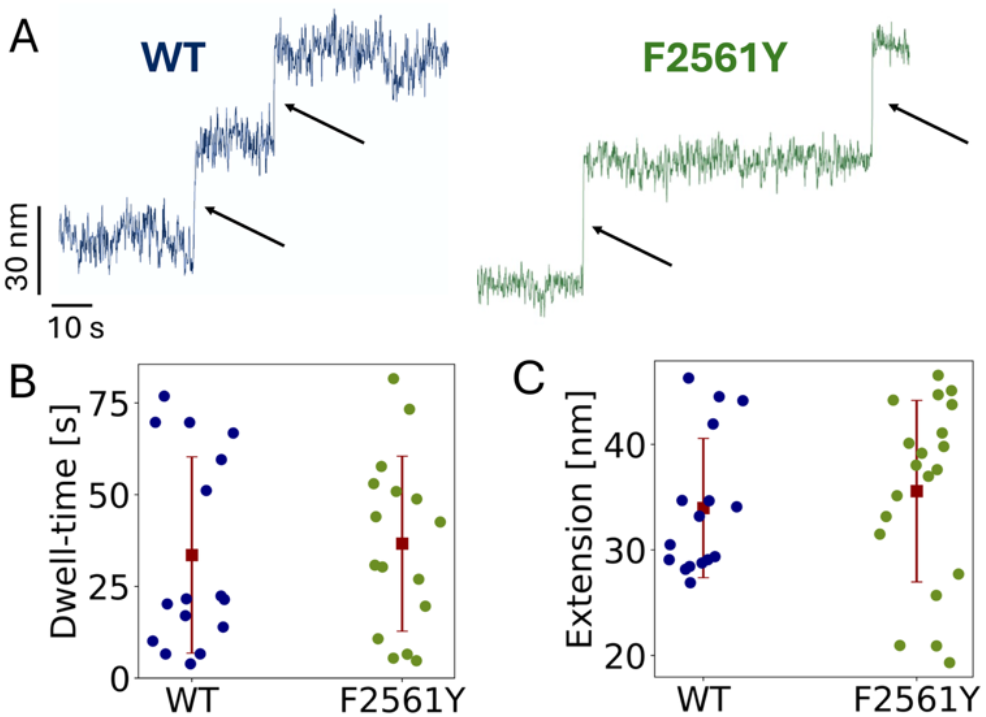
Force-induced unfolding of A2 domains in wildtype and F2561Y VWF dimers. (A) Representative extension-time traces showing two sequential A2 domain unfolding events in a VWF dimer for wildtype (blue) and the F2561Y mutant (green). Arrows point to the individual unfolding events. B) Dwell-times observed at 12.5 pN for A2 domain unfolding in wildtype (blue; WT: 33.6 ± 26.7 s) and F2561Y (green; F2561Y: 36.7 ± 23.9 s) VWF dimers show no significant difference (Welch’s *t*-test: *p* = 0.71). C) Comparison of extension steps associated with A2 domain unfolding for wildtype (WT: 32.5 ± 7.7 nm) and F2561Y (F2561Y 35.7 ± 8.6 nm) VWF show no significant difference (Welch’s *t*-test: *p* = 0.24). Data points in panels B and C are from individual measurements on VWF dimers; red squares and error bars are the mean ± standard deviation of all values for the given condition.

We occasionally observed additional, larger extension steps within the same force range as the A2 unfolding events, which we attribute to dissociation of the D4-D4 domain interaction in the dimer (Figure 1F, Supplementary Figure S3). Consistent with previous MT observations [9], we find that this dissociation can occasionally be reversible. For mutant and wildtype, the extension increases attributed to the dissociation of the D4-D4 interactions (defined as steps ≥ 50 nm) are within experimental error (WT: 96.3 ± 28 nm; F2561Y: 96 ± 27.5 nm) and substantially larger than the length increases arising from A2 domain unfolding, yet occur in a similar force range (14 pN ± 7 pN for wt and 13 pN ± 2 pN for the F2561Y mutant; Supplementary Figure S3).

Previous AFM force spectroscopy experiments reported D4-D4 dissociation at much higher forces (≈ 50 pN), under conditions of higher loading rates [10]. In contrast, magnetic tweezers measurements operate at constant forces, corresponding to (close to) zero loading rate. The observation that D4-D4 dissociation occurs at much lower forces in the MT assay compared to AFM force spectroscopy suggests a strong loading-rate dependence of the D4-D4 interaction, larger than for A2 unfolding. A strong loading rate dependence implies that the D4-D4 interaction involves a longer-range energy barrier with a larger distance to the molecular transitions state.

### The F2561Y Mutation Shifts VWF Stem Opening to Lower Forces

Using the A2 unfolding steps as a molecular fingerprint to identify properly attached VWF dimers, we next investigated the behavior of VWF dimers at low forces, where the force response is dominated by the VWF stem (9). To assess the impact of F2561Y on stem mechanics, we monitored force-induced stem opening and closing transitions in VWF dimers at low forces using magnetic tweezers (Figure 3). These transitions occur after dissociation of the D4-D4 interaction and manifest as stochastic switching between compact (closed) and extended (open) stem conformations (Figure 3A,B,C). Representative extension-time traces recorded at several forces reveal reversible transitions for both wildtype and mutant dimers. Higher extensions correspond to the open stem conformation, whereas lower extensions reflect more compact states. As the applied force decreases, the stem adopts more closed and compact conformations, resulting in traces dominated by lower extensions. For the wildtype an approximately equal population of open and closed conformations is observed at 1 pN (Figure 3B), indicating the equilibrium force for this transition. In contrast, the F2671Y mutant reaches this equilibrium at lower forces (Figure 3C), and the fully closed stem conformation is only rarely observed.

**Figure 3:**
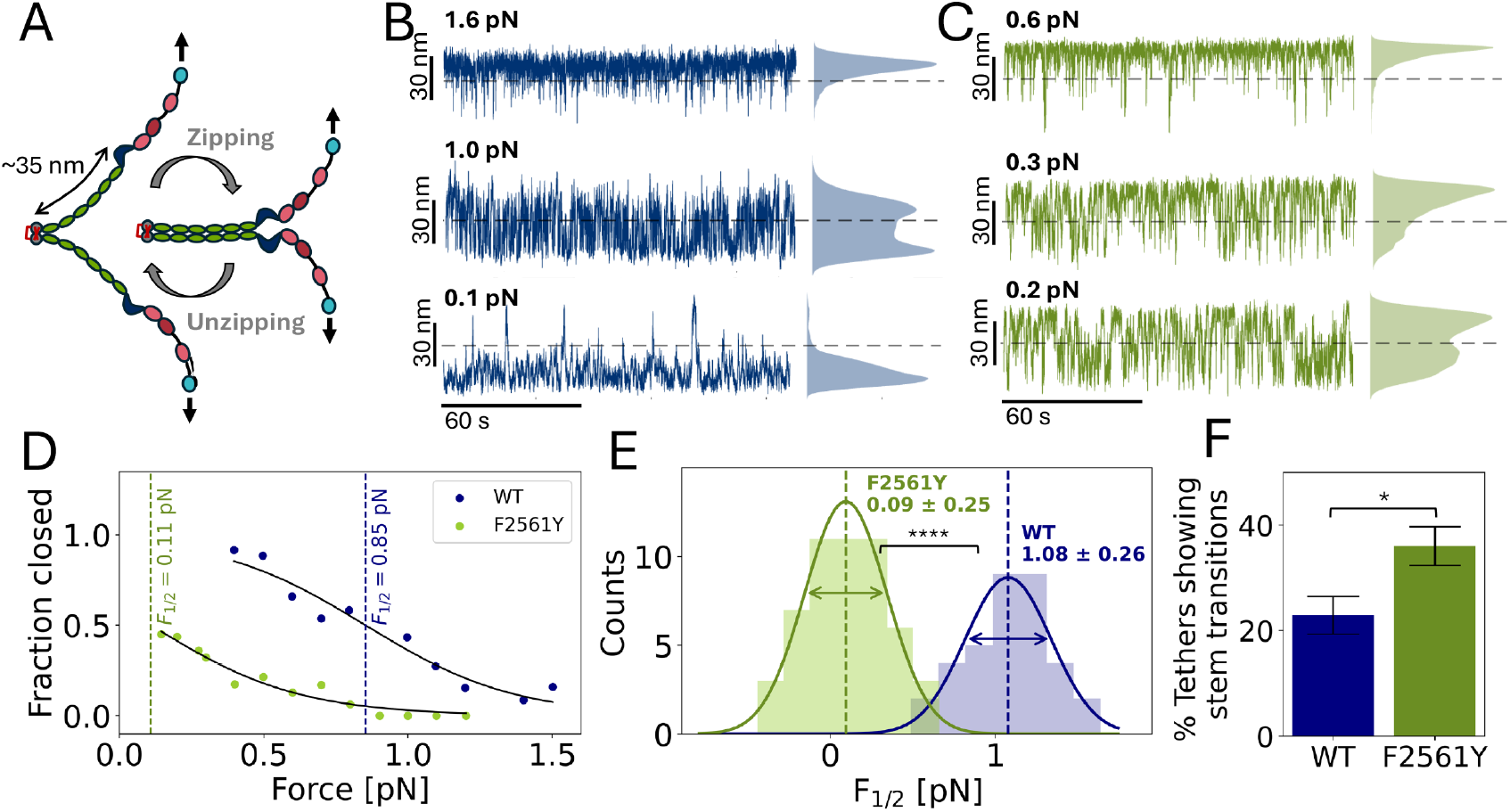
The F2561Y mutation shifts VWF stem opening to lower forces. A) Schematic illustration of reversible opening and closing of the VWF stem in a zipper-like manner. Arrows indicate the direction of force application in the magnetics tweezers. B, C) Representative extension-time traces and corresponding kernel density estimates (to the right of the traces) obtained at low forces for wildtype (B) and the F2671Y mutant (C). Dashed lines indicate the midpoint extension. D) Fraction of closed stem configuration as a function of applied force for two representative molecules of WT VWF (blue) and the F2561Y mutant (green). Solid lines are fits to the two-state model (Equation 1). Vertical lines denote the fitted midpoint forces F_1/2_. E) Distributions of midpoint forces F_1/2_ for WT (blue) and F2561Y (green) VWF dimers (N_WT_ = 35, N_F2561Y_ = 52). Solid lines are Gaussian fits. Vertical lines visualize the mean F_1/2_ values and arrows denote the standard deviation. The midpoint force is significantly reduced for F2561Y compared to wildtype (WT: 1.08 ±0.26 pN, F2561Y: 0.09 ±0.25 pN; Welch’s *t*-test, *p* < 0.0001). F) Fraction of VWF dimers that exhibit stem transitions. The fraction is significantly higher for the F2561Y mutant (green, 36% ± 3.69%, N = 169) than for the wildtype (WT, blue, 22.9% ± 3.55%, N = 140) (χ^2^-test, *p* = 0.013). Dimers that do not show stem transitions likely exhibit a D4-D4 interaction, which holds the stem closed.

To quantitively analyze these transitions, we employ a simple two-state model. We define a midpoint extension threshold and use it to classify the extensions into a closed and an open state. We then analyze the extension time traces by computing the fraction of closed stem conformation as a function of applied force, from which we obtain the midpoint force F_1/2_ from a fit of the two-state model presented in Equation 1 (Figure 3D). We find average midpoint forces of F_1/2_ = 1.08 ±0.26 pN (N = 35) for the wildtype, which is consistent with previous measurements that reported F_1/2_ = 1.00 ±0.24 pN [9], and F_1/2_ = 0.09 ±0.25 pN (N = 52) for the F2561Y mutant (Figure 3E). The shift toward lower midpoint forces in the mutant by about one order of magnitude is highly statistically significant (Welch’s *t*-test, *p* < 0.0001).

In addition, a significantly larger fraction of tethered F2561Y dimers (36 % ± 3.69%) exhibited detectable stem opening and closing compared to wildtype (22.9 % ± 3.55%) (χ^2^-test, *p* = 0.013; Figure 3F). The increased occurrence of stem transitions suggests an altered conformational equilibrium in the mutant that favors dynamic stem behavior within the probed force range, indicating less stabilization of the compacted stem conformation. Together, these findings demonstrate that the F2561Y mutation destabilizes stem compaction, shifting stem opening to substantially lower forces and thereby increasing the mechanical activation propensity of VWF.

### AFM Images Reveal a Mainly Open Stem Conformation of VWF F2561Y

To obtain structural evidence complementary to the single-molecule force spectroscopy measurements, we analyzed VWF dimers by AFM imaging. Representative AFM images of wildtype and F2561Y dimers deposited on mica and imaged in air reveal distinct differences in stem morphology (Figure 4A,B). Individual molecules were classified according to their stem configuration as compact (closed) when they displayed three distinct protrusions or as extended (open) when they exhibited two longer flanks (Figure 3C; see Methods for details). While both conformations were observed for wildtype and mutant proteins, open stem configurations occurred markedly more frequently for the F2561Y variant. Quantitative analysis of a large number of molecules (N_wt_ = 69, N_F2561Y_ = 167) shows a significant increase in the fraction of dimers exhibiting an open stem in the F2561Y mutant (76% ± 9%) compared to the wildtype (39% ± 8%) (Figure 4D; χ^2^ test, *p* < 0.0001). The fraction of compact wildtype dimers is in good agreement with previous measurements obtained under identical buffer conditions (pH 7.4, in the presence of divalent ions), representative of physiological plasma conditions [10, 52].

**Figure 4:**
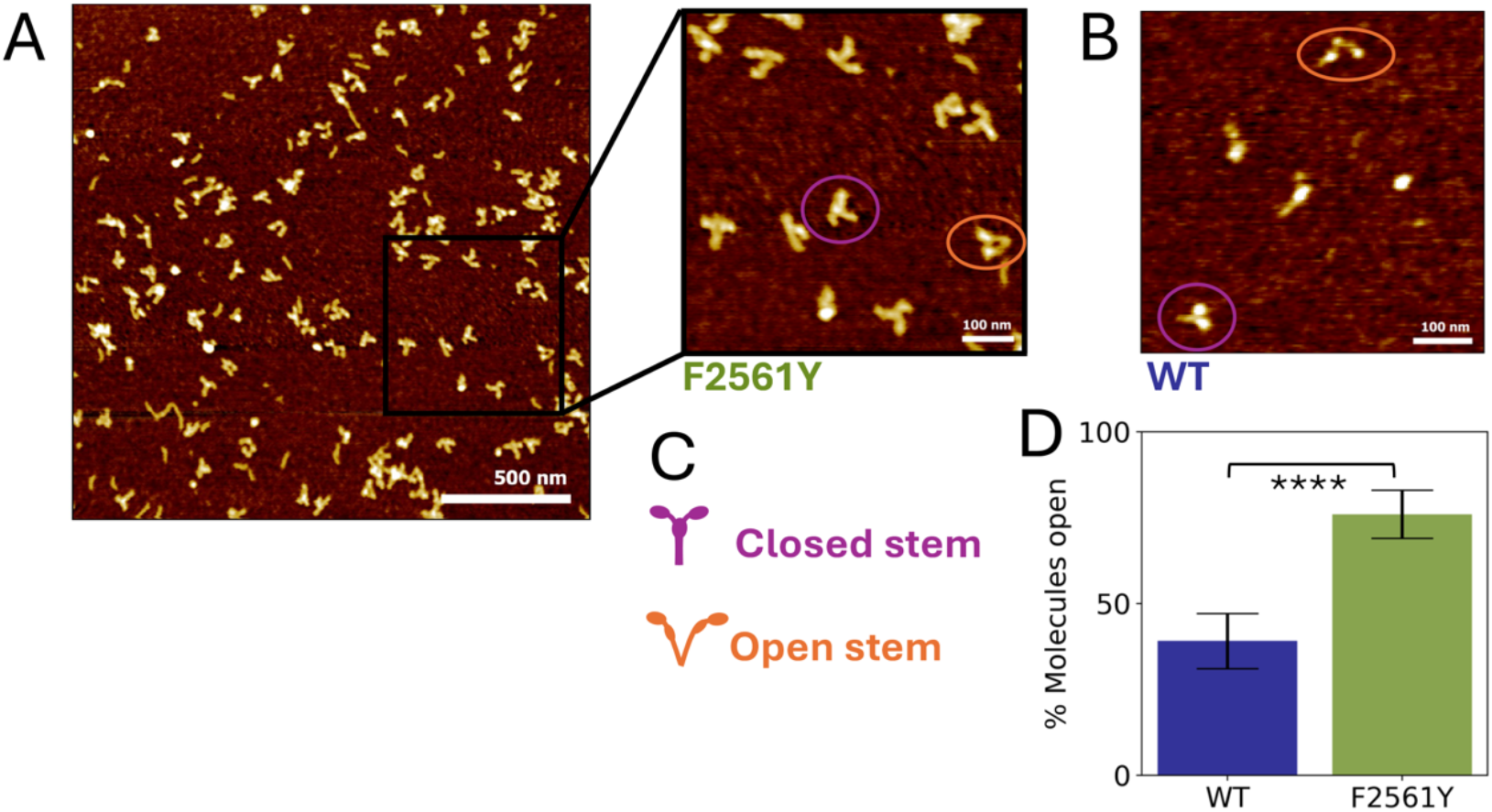
The F2561Y mutation increases the fraction of VWF dimers with an open stem conformation. A, B) Representative AFM images of VWF dimers deposited on mica and imaged in air. Panel A shows the F2561Y mutant (overview and zoomed-in view), and panel B shows a zoomed-in image of the wildtype VWF. Purple and orange circles mark examples of dimers identified with closed and open stem configuration, correspondingly. C) Simplified scheme for closed (purple) and open (orange) VWF dimers. Spheres represent folded domains, corresponding to brighter, and thus higher features in the AFM image. D) Quantification of VWF dimers exhibiting an open stem configuration, determined from AFM images. The F2561Y mutant (green; 76 ± 9%) shows a significantly higher fraction of open molecules compared to the wildtype (blue; 39 ± 8%) (χ^2^ test, *p* < 0.0001).

These structural data independently confirm the single-molecule force spectroscopy results, which demonstrate a shift in the conformational distribution of the VWF stem toward more extended states in the mutant. The increased prevalence of open conformations observed by AFM is therefore consistent with a destabilization of the compact stem state and a redistribution of the conformational equilibrium toward extended configurations.

## Discussion

In this study, we investigate how the prothrombotic VWF F2561Y mutation alters mechanical regulation of the VWF stem using single-molecule magnetic tweezers complemented by AFM imaging. While the mutation does not affect unfolding of the A2 domains, it profoundly alters the force-dependent behavior of the VWF stem, revealing a stem-specific mechanical phenotype.

Unfolding of the A2 domain, which is a key regulatory transition controlling multimer size by ADAMTS13 cleavage *in vivo*, serves as an internal mechanical fingerprint of correctly tethered VWF dimers in our assay. We observe that A2 unfolding contour lengths, unfolding rates, and force responses are indistinguishable between wildtype and F2561Y VWF dimers, which suggests that the mutation in the C4 domain does not globally destabilize VWF. Instead, the mechanical effects of F2561Y are spatially confined, pointing toward localized perturbation within the stem region.

In contrast to A2 behavior, stem mechanics are strongly altered by the F2561Y mutation. Magnetic tweezers measurements reveal that stem opening transitions occur at markedly lower forces in the F2561Y mutant. The midpoint force for stem opening is shifted from ~1.1 pN in wildtype to ~ 0.1 pN in the mutant, demonstrating that substantially less mechanical load is required to access extended stem conformations. Moreover, a larger fraction of mutant molecules exhibits stem transitions within the probed force range. Together, these findings indicate that the compact stem conformation is mechanically destabilized in F2561Y VWF.

AFM imaging provides independent structural support for this conclusion. Analysis of a large population of surface-deposited VWF dimers reveals a statistically significant enrichment of open stem conformations in the mutant. Although AFM images represent a static snapshot in the absence of applied force, the observed shift in conformational distribution is fully consistent with the reduced mechanical stability of the compact stem observed in magnetic tweezers experiments.

The F2561Y mutation is located in the C4 domain, a component of the dimeric stem that also harbors the RGD motif for platelet integrin GPIIb/IIa binding. Previous structural and spectroscopic studies reported preserved folding of isolated C-domains but altered spectral features in dimeric constructs [26, 31], suggesting that F2561Y perturbs interdomain organization rather than intrinsic domain stability. Our data support these findings and further provide direct mechanical observations, indicating that the mutation introduces a subtle structural perturbation at C-domain interfaces that does not alter overall stem architecture and A2 mechanosensing, but weakens the packing interactions that stabilize a compact stem conformation.

Within the current structural model of VWF, compaction of the stem is reinforced by intersubunit D4-D4 interaction, which acts as a mechanical clamp that enables zipper-like stem closure. The observed increased population of VWF F2561Y molecules showing stem transitions in magnetic tweezers experiments and extended conformations in AFM imaging are consistent with a reduced effective stabilization of this D4-D4 interaction or a decreased propensity of the stem to re-form the compact state. In either scenario, the conformational equilibrium is shifted toward more extended configurations.

This altered mechanical landscape provides a direct link to the gain-of-function phenotype of the F2561Y variant. A stem that opens more readily under low hydrodynamic forces would increase effective contour length and enhance accessibility of platelet-binding sites within the C4 domain, thereby lowering the force threshold for VWF activation. This mechanism is consistent with previous reports of increased shear sensitivity and enhanced platelet aggregate formation [26, 27] and provides a molecular explanation for the prothrombotic phenotype associated with F2561Y.

The combination of high-resolution magnetic tweezers, which is uniquely suited to probe low-force conformational equilibria such as VWF stem opening in the sub-piconewton regime over long observation times, and AFM imaging of a large molecular population provides complementary mechanical and structural evidence for altered conformational distributions. Together these approaches enable a quantitative view of how a single point mutation reshapes the mechanical energy landscape of VWF. Beyond the role of the F2561Y mutation, this work establishes magnetic tweezers as a powerful platform for probing conformational transitions in the very low-force regime and for quantitatively linking molecular mechanics to hemostatic function.

## Acknowledgement

We thank Adina Hausch, Sofia Gruber, and Baris Yoldas Cinemre for help with initial MT measurements and discussions, Thomas Nicolaus, Dave van den Heuvel, Elleke van Harten, Maria Kurilova, Sidonie Lieber, Andreas Spörhase, and Olga Ustinov for laboratory support, and Gerhard Blab, Reinhard Schneppenheim, Tor Sewring, Alptuğ Ulugöl, and Matthias Wilmanns for useful discussions. This work was supported by Utrecht University, by the European Research Council Consolidator Grant “ProForce” (101002656), and a Feodor-Lynen fellowship from the Alexander von Humboldt Foundation.

## Authorship Contributions

Contribution: C.K., M.B., and J.L. designed research; C.K. and S.D.P. performed experiments; C.K. and T.O. expressed proteins; C.K. and S.D.P. analyzed experimental data; and C.K. and J.L. wrote the paper with input from all authors.

