## Supplementary Information for "Magnetic Tweezers Experiments Reveal Increased Mechanical Sensitivity of the F2561Y Mutant in Von Willebrand Factor"

**Content:**

**Supplementary Materials and Methods**
**Supplementary Table S1**
**Supplementary Figure S1-S3**

### Supplementary Materials and Methods

#### ELP Linkers

Elastin-like polypeptide linkers are used for protein attachment and as a spacer between the flow cell surface and the VWF construct. Recombinant ELP linkers with a contour length of ~120 nm consist of ~300 amino acids with the sequence $[(VPGEG)\text{-}(VPGVG)_4\text{-}(VPGAG)_2\text{-}(VPGGG)_2\text{-}(VPGEG)]_6$, bearing a single N-terminal cysteine residue for linkage to maleimide groups on the glass surface, and a C-terminal *LPETGG* recognition motif for sortase-mediated protein attachment.
Cloning and expression was performed following standard procedures as previously described (1, 2), and the linkers purified with the ITC method, a purification method based on the thermal response of the ELPs with repeated thermoprecipitation at 60 °C followed by redissolution at 4 °C (3).

#### Enzyme Expression and Purification

The enzymes used for protein attachment in the flow cells, i.e. sfp-phosphopantetheinyl transferase (2) for protein biotinylation and eSortaseA (4, 5) for linkage to ELP-linkers, were expressed in *E. coli* DE3 cells and purified following standard protocols. Bacteria were cultured in LB medium (Carl Roth), supplemented with kanamycin or ampicillin, inducted with IPTG (Sigma-Aldrich), and proteins expressed at 18 °C for ~24 h. Cell pellets were lysed by incubating in lysis buffer (Supplementary Table S1) on a rolling incubator, followed by tip-sonication. After filtering to 0.22 µm, the supernatant was purified over a Ni-sepharose HisTrap FF column (5 ml, Citiva) with the appropriate binding and eluted with additional 300 mM imidazole. Purified fractions were verified in an SDS gel and further concentrated in the corresponding storage buffer (Supplementary Table S1) using a 3 kDa Amicon centrifugal filter (Merck). Concentrated proteins were aliquoted for direct usage and stored at -80 °C.

#### MT Setup

The MT measurements used a custom-built MT setup as previously described (6-8). A pair of vertical oriented 5x5x5 $mm^3$ permanent magnets (W-05-N50-G, Supermagnete) is placed above the flow cell in an external magnet holder with a gap of 1 mm (9). The height of the magnets is controlled by a DC-motor (M-126.PD2; Physik Instrumente) with mercury controller. The beads are illuminated with an LED source (650 nm, Thorlabs) and imaged using a 40x oil immersion objective (UPLFLN 40x, Olympus), placed on a piezo stage (Pifoc P-726.1CD, Physik Instrumente) which can be moved to adjust the focus and record a z-look-up-table for tracking of the bead’s diffraction rings (10). The field of view of 680x680 $\mu m^2$ is imaged using a CMOS sensor camera (CP80-25-M-72, Optronis) with 5120x5120 pixels, operating at a sampling frequency of 72 Hz. Images are transferred to a frame grabber (microEnable

5 ironman VQ8-CXP6D, Silicon Software) and analyzed with a custom-written open-source LabView code (11). Forces were calibrated based on transverse fluctuations of 21 kbp long double-stranded DNA tethers (9, 12), using a tracking frequency of 400 Hz in a reduced field of view and the Hadamard variance as analysis method (Supplementary Figure S1) (8, 13, 14).

### Supplementary Table

**Supplementary Table S1:** Buffers Magnetic Tweezers measurements and AFM imaging, and for protein purification from a Ni-sepharose HisTrap FF column for eSortase A and sfp-phosphopantetheinyl. All buffers are filter-sterilized using a nitrocellulose filter membrane with a pore size of 0.22 µm (Merck Millipore); HPLC buffers are additionally degassed.

| **Buffer** | **Components** |
|---|---|
| VWF measurement buffer | 20 mM Hepes, 150 mM NaCl, 1 mM $CaCl_2$, 1 mM $MgCl_2$,<br>pH 7.4 |
| Cell lysis buffer | 50 mM Tris-HCL, 50 mM NaCl, 5 mM $MgCl_2$, 0.1% (v/v) TritonX-100, 10% (v/v) glycerol,<br>pH 7.5,<br>DNAse I (Merck), Lysozyme (Merck), EDTA-free cOmplete protease inhibitor (Merck) |
| eSortase A binding buffer | 25 mM Tris-base, 300 mM NaCl, 20 mM Imidazole, 0.25% (v/v) Tween-20, 10% (v/v) glycerol,<br>pH 7.8 |
| Sfp phosphopantetheinyl transferase binding buffer | 20 mM Tris-HCl, 500 mM NaCl, 5 mM imidazole,<br>pH 7.5 |
| eSortase A storage buffer | 25 mM Tris-base, 75 mM NaCl, 1 mM $CaCl_2$,<br>pH 7.2 |
| Sfp phosphopantetheinyl transferase storage buffer | 50 mM Hepes, 150 mM NaCl, 10% (v/v) glycerol,<br>pH 7.5 |

#### Supplementary Figures

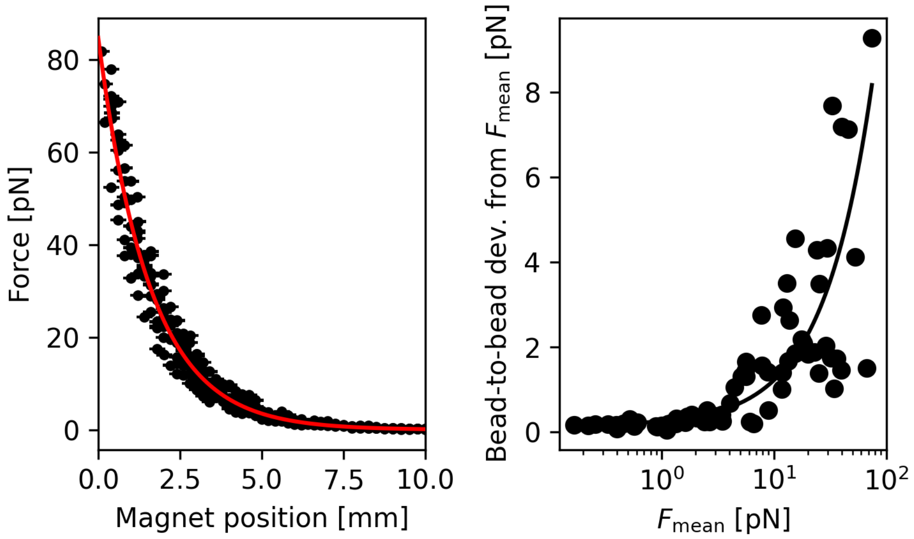


**Supplementary Figure S1: Force calibration curve of the magnetic tweezers setup with M270 magnetic beads.** Left: Force over magnet position for 21 kbp long dsDNA (9, 12). Forces are calculated from the transverse bead fluctuations using the Hadamard variance method (8, 13), with errorbars representing the standard error ($\sigma$). From the exponential fit ($F(x) = F_{max}e^{-x/\lambda}$, red) we derive a maximum force $F_{max}$ of 84.7 pN and a decay length $\lambda$ of 1.6 mm. Right: The bead-to-bead deviations from $F_{mean}$, i.e. $\sigma_{F_{mean}}$, are fitted using $\sigma_{F_{mean}} = \sigma_0 + \sigma_{rel}\, F$, with $\sigma_{rel}$ the relative (unitless) deviation, and $\sigma_0$ (pN) accounts for intrinsic system errors. The fitted values are $\sigma_{rel} = 10.8\%$ and $\sigma_0 = 0.14$ pN.

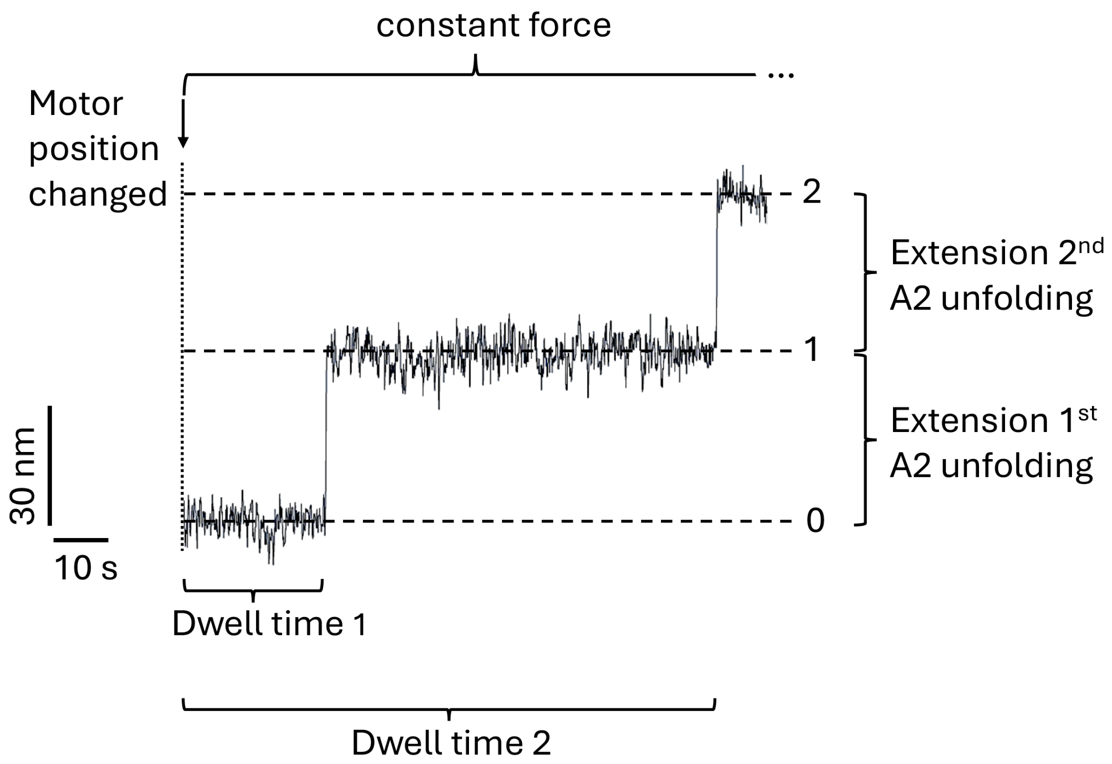


**Supplementary Figure S2: Dwell time and extension analysis of A2 domain unfoldings.** A representative extension-time trace of a VWF dimer in magnetic tweezers recorded at a constant force of 12.5 pN is used to visualize the analysis of A2 domain unfolding events. The dashed lines indicate the mean extension at a distinct step: ‘0’ shows the lowest extension with both A2 domains folded, ‘1’ indicates the mean extension of the dimer with one A2 domain unfolded, and ‘2’ is the dimer with both A2 domains unfolded. This pattern is used as a unique fingerprint in our measurements to identify a single intact and correctly attached VWF dimer. A2 unfolding is reversible and can be observed several times in the same molecule. The analyzed extensions used for Figure 2C correspond to the length increments ‘0’ → ‘1’ and ‘1’ → ‘2’. Dwell times are analyzed from the start of the constant force plateau, i.e. the time point when the magnet is moved to a distinct position. ‘Dwell time 1’ is the measured time from the start of the force plateau until the first distinct step in extension of ~ 40 nm is observable, which corresponds to the unfolding of one A2 domain. ‘Dwell time 2’ is measured exactly like ‘Dwell time 1’, with the starting time being the start of the new force plateau, until the second distinct step in extension of ~ 40 nm, which corresponds to the unfolding of the second A2 domain. Since both A2 domains unfold independently, the dwell times are considered independent and are therefore both measured from the start of the force plateau, i.e. ‘Dwell time 2’ is measured from the beginning of the measurement at the corresponding force (and not measured from the first to the second A2 domain unfolding).

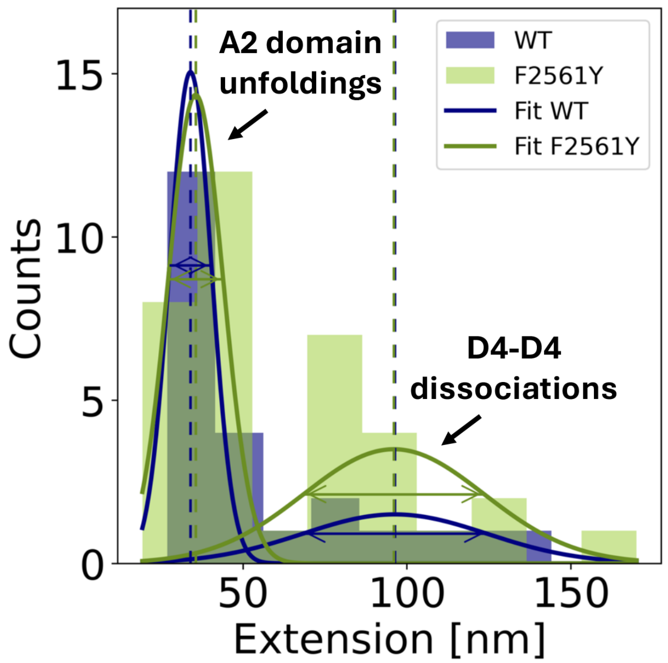


**Supplementary Figure S3: Distribution of extension steps in VWF dimers.** Histogram of all observed extension steps for wildtype (blue) and F2561Y (green) dimers, observed in magnetic tweezers measurements. Two distinct populations of extension events are observable, associated with A2 domain unfolding and D4-D4 dissociation (7). A threshold (50 nm) was determined by minimizing the total weighted variance of two Gaussian-distributed populations across all measured extension steps. Gaussian fits to the separated populations yielded mean extension values of 34 ± 6.4 nm (A2) and 96.3 ± 28 nm (D4-D4) for wildtype and 35.6 ± 8.4 nm (A2) and 96 ± 27.5 nm (D4-D4) for F2561Y. For either population, no significant difference wa observed between wildtype and F2671Y ( Welch's *t*-test: A2: $p = 0.54$; D4: $p = 0.98$). The A2 unfoldings serve as a molecular fingerprint to identify correctly attached dimers. The dissociation of the D4-D4 interaction is not happening in all traces.
